# Unraveling the Time-Frequency Features of Emotional Regulation: Based on an Interpretable XGBoost-SHAP Analytical Framework

**DOI:** 10.1101/2024.03.15.585273

**Authors:** Cheng Si

## Abstract

Negative emotions, while crucial for survival, can lead to adverse health effects if not managed properly. Our understanding of temporal EEG changes during emotion regulation is limited. To address this gap, this study employs interpretable machine learning techniques, XGBoost-SHAP model, to analyze EEG data. This study investigates the neural mechanisms underlying emotion regulation, with a focus on EEG oscillations in the lateral prefrontal area channels (F3, F4, F7, F8) across four specific frequency bands (Alpha, Beta, Theta, Delta). By identifying predictive features and patterns, this approach offers insights into the temporal dynamics of emotion regulation and the involvement of specific brain regions, enhancing our understanding of emotional processing and providing avenues for effective interventions. The findings reveal a significant relationship between specific EEG feature changes and emotional ratings during the emotion regulation process. The LPFC emerges as central in cognitive control and emotional regulation. These results highlight the LPFC’s rapid and effective role in regulating complex emotional dynamics, crucial for understanding and treating emotional disorders. The study underscores the importance of machine learning in elucidating neural mechanisms and guiding personalized interventions for emotional well-being.

## Introduction

Negative emotions play a crucial role in human survival and adaptation, offering evolutionary advantages. However, inadequate management of negative emotions can lead to detrimental health effects in the long term, including cardiovascular diseases, depression, and anxiety (1),(2),(3),(4),(5). Emotion regulation encompasses the process of modulating the type, timing, and expression of emotions (6). It holds a pivotal position in human survival, serving as a defense mechanism against emotional dysfunction and contributing significantly to the maintenance of social harmony and stability (7).

Extensive research has indicated the critical involvement of the lateral prefrontal cortex (LPFC) in emotional experience and regulation, including ventrolateral prefrontal cortex (VLPFC) and dorsolateral prefrontal cortex (DLPFC) (8),(9),(10),(11),(12),(13). Broadly, the DLPFC is associated with conflict monitoring and attention maintenance, while the VLPFC is implicated in semantic organization and language processing (14), 15), 16).

Numerous cognitive processes, such as learning, memory, decision-making, and perception, involve dynamic information processing and integration. Investigating the brain’s dynamic features aids in understanding how it processes and integrates information across different temporal points, shedding light on cognitive processes and collaborative brain region interactions. Gross expanded his early model of emotion regulation processes with the Expanded Process Model (EPM), which places temporal dynamics at its core (17). In the EPM, initial emotion perception activates and identifies emotion regulation goals, subsequently leading to goal execution. Successful strategy implementation modifies primary appraisal system features, sustaining emotion regulation effects until goal deactivation. From a cognitive dynamic theory perspective, emotion regulation unfolds as a dynamic process to meet regulation goals, rather than statically maintained within specific brain areas. Some researchers have examined cognitive reappraisal’s temporal processes through EEG oscillations. Ertl et al. (2013) found significant differences in emotion downregulation compared to maintenance regulation in the Theta (4-8 Hz) band energy at 1-5 s, possibly linked to Theta waves’ role in regulating emotional responses and promoting stability (18). However, our understanding of EEG frequency changes related to emotion regulation over time remains limited.

XGBoost stands as a potent machine learning algorithm, whose interpretability advantage makes it an ideal choice for investigating dynamic processes in EEG data. Firstly, interpretability is a crucial factor in ensuring the credibility of research findings. In the field of neuroscience, understanding the decision-making process of models is vital for comprehending the mechanisms underlying brain activity (19), 20). The XGBoost model provides a clear means of discerning key features and patterns influencing emotion regulation within EEG data (21). Secondly, the interpretability of the XGBoost model aids in constructing more persuasive theoretical models, thus deepening our understanding of the dynamic processes of emotion regulation. By identifying specific EEG patterns associated with successful emotion regulation, we can provide a more detailed and accurate description of the neural basis of emotion regulation. Lastly, employing interpretable machine learning models can facilitate the translation and application of research findings.

The implications of this research extend beyond the academic sphere, offering potential applications in clinical practice and the development of personalized cognitive enhancement tools. By leveraging machine learning to elucidate the neural underpinnings of emotional regulation, this study contributes to a deeper understanding of the mechanisms by which individuals modulate their emotional responses. Such knowledge is crucial for advancing our grasp of emotional disorders and refining therapeutic strategies aimed at fostering healthier emotional processing and regulation.

## Methodology

### Data Source

The data for this analysis were sourced from Li et al. (2022), incorporating the data of 26 participants who received vertex stimulation (to avoid interference from TMS stimulation of the VLPFC). All participants were right-handed, with no history of neurological diseases or brain injuries, healthy at the time of the experiment, and with normal or corrected-to-normal vision. Prior to the experiment, informed consent was obtained, and participants were compensated with 100 RMB upon completion. The data include conditions across 3 (social feedback valence: positive, neutral, and negative) × 2 (regulation type: viewing and reappraisal), totaling 6 conditions. The experiment utilized 60 identity photos of young individuals with neutral facial expressions (30 males and 30 females), with all photos standardized in background color (white), resolution, and brightness. According to ratings from another homogeneous participant group, there were no significant statistical differences in attractiveness and supportiveness scores among the six groups of pictures (*F*s < 1). For specific experimental procedures, refer to the original paper (22).

### EEG Preprocessing and Feature Extraction

Raw data were collected via a 32-channel amplifier (NeuSen.W32, Neuracle) at a sampling rate of 250 Hz, with electrode impedance kept below 10 kΩ. The reference electrode was placed at TP9, without the application of online filters. Initially, the data underwent a bipolar mastoid reference (TP9 TP10), followed by band-pass filtering from 0.1-30 Hz and notch filtering around 50 Hz to remove the influence of electrical noise. The filtered data were segmented from 500 ms before to 4000 ms after picture presentation, resulting in segments of 4500 ms length, and baseline corrected using the -500 to 0 ms data. The segmented data were then subjected to fast ICA for independent component analysis, with eye movement and muscle activity-related ICA components automatically excluded. The cleaned data underwent Morlet wavelet transformation, yielding energy values for the Delta, Theta, Alpha, and Beta frequency bands. This study focused primarily on the four electrode channels near the DLPFC and VLPFC, namely F3, F4, F7, F8. For each participant, energy in these channels across four segments was averaged in 100 ms steps, resulting in a dataset of 26 (participants) × 90 (pictures) × 2 (conditions: viewing, reappraisal), totaling 4680 segments, with 40 (time segments) × 4 (frequency bands), equating to 640 features. The dataset for machine learning classification consisted of 4680 × 640 data points.

### Model Optimization

The XGBoost is an efficient classification machine learning model. This study employed the XGBoost Classifier to classify viewing and reappraisal conditions. Due to the plethora of hyperparameters in XGBoost, this research initially defined a hyperparameter space and conducted 500 random trials across approximately 18,000 parameter combinations (hyperparameter space detailed in 1). The model parameters that achieved the highest AUC were selected for the final model. The relevant code for building the machine learning model has been uploaded to GitHub (https://github.com/Cheng990427/Paper-1).

### Model Validation

In our experiments, the dataset was split into a training set and a testing/validation set at a ratio of 80% and 20%, respectively. For the validation set, we utilized a ten-fold cross-validation method for verification. The results indicated that our model achieved an average accuracy of 69% on the validation set. For each subject, the ten-fold cross-validation results for individual subject prediction accuracy ranged from 0.63 to 0.98. Additionally, the model was evaluated on multiple performance metrics on the XG-Boost model. The model demonstrated good performance in terms of AUC, accuracy, sensitivity, precision, and F1 score, with the specific results as follows: the AUC was 0.76, with a 95% confidence interval of [0.75, 0.78]; the accuracy was 0.69, with a 95% confidence interval of [0.68, 0.71]; the sensitivity was 0.68, with a 95% confidence interval of [0.67, 0.70]; the precision was 0.69, with a 95% confidence interval of [0.68, 0.71]; the F1 score was 0.69, with a 95% confidence interval of [0.67, 0.70]. These results demonstrate that the model possesses acceptable accuracy and stability, as seen in Table 2. The aforementioned results highlight the good performance of our model in the given task.

**Table 1.** XGBoost Hyperparameter Space.

| Hyperparameter | Hyperparameter Space (start, stop, step) |
| --- | --- |
| objective | binary:logistic |
| booster | gbtree |
| learning_rate | 0.05, 0.30, 0.05 |
| max_depth | 1, 49, 1 |
| n_estimators | 100, 1100, 100 |
| min_child_weight | 3, 8, 1 |
| subsample | 0.4, 1, 0.1 |
| seed | 1444 |

**Table 2.** Important Features for Predicting Emotional Regulation and Passive Viewing.

| No. | Positive Predictive Feature Index | Negative Predictive Feature Index |
| --- | --- | --- |
| 1 | F3-0-Alpha | F3-600-Beta |
| 2 | F3-900-Alpha | F3-700-Beta |
| 3 | F3-1000-Alpha | F3-800-Beta |
| 4 | F3-1100-Alpha | F4-3200-Delta |
| 5 | F3-1200-Alpha | F4-1700-Theta |
| 6 | F3-1300-Alpha | F4-2700-Theta |
| 7 | F3-3000-Alpha | F4-900-Beta |
| 8 | F3-3300-Beta | F7-200-Delta |
| 9 | F3-3400-Beta | F7-3500-Delta |
| 10 | F3-3500-Beta | F7-100-Theta |
| 11 | F4-1600-Theta | F7-500-Theta |
| 12 | F4-2800-Theta | F7-1900-Theta |
| 13 | F4-3900-Alpha | F7-2000-Theta |
| 14 | F4-1600-Beta | F7-3200-Theta |
| 15 | F4-2100-Beta | F7-3200-Beta |
| 16 | F7-1000-Theta | F8-100-Delta |
| 17 | F8-3800-Delta | F8-1000-Alpha |
| 18 | F8-500-Alpha | F8-1300-Alpha |
| 19 | F8-600-Alpha | F8-2300-Beta |
| 20 | F8-3300-Alpha | F8-3000-Beta |

### Model Explanation

The SHAP (Shapley Additive Explanations) interpreter adopts the concept of Shapley values from game theory as an advanced tool for elucidation, aimed at clarifying the predictive behavior of machine learning models (23). Originally applied in cooperative game scenarios, Shapley values quantify the individual contributions of participants to the collective outcome. Within the SHAP framework, input features of a model are analogized to players in a game, with the model’s predictive output equating to the total payoff. To compute a feature’s SHAP value, the model considers the role of this feature across all possible subsets of features. For each potential subset of features, the change in model prediction with and without the target feature is calculated, representing the marginal contribution. Averaging the marginal contributions of a feature across all possible combinations yields the Shapley value for that feature. For a given single prediction, the sum of SHAP values for all features, plus a baseline prediction value (usually the average prediction over the training data), should equal the actual model prediction for that sample.

Utilizing SHAP values enables the quantification of each feature’s contribution to the model’s prediction outcome, thereby decoding the significance of features in the model’s decision-making process. This method’s versatility makes it applicable to a variety of machine learning models, including linear models, neural networks, and random forests, hence its widespread use in the field of model interpretability. In this study, by calculating and ranking the SHAP values of all features, we further filtered out the top 40 features by | SHAP value | as significant features. These prominent features played a crucial role in the model’s classification of watching behavior versus re-rating behavior, significantly surpassing other features in influence. Through this method, we can identify key predictive factors within the model and deepen our understanding of the model’s decision-making logic.

## Results

### Important Feature Selection Based on SHAP Values

The original SHAP values for each feature were obtained using the SHAP interpreter, and the importance ranking of all features was determined based on the absolute magnitude of the SHAP values. Figure1 A displays the top 20 features ranked by their original SHAP values, while Figure1 B illustrates the importance of features during each random sampling process. Subsequently, the top 20 features with positive and negative impacts were selected based on the ranking of | SHAP value |, resulting in a total of 40 important features identified in this model. Table2 lists the top 20 positive predictive features and the top 20 negative predictive features. Figure1 C presents the distribution of these 40 important features across four frequency bands and all time points.

**Fig. 1.**
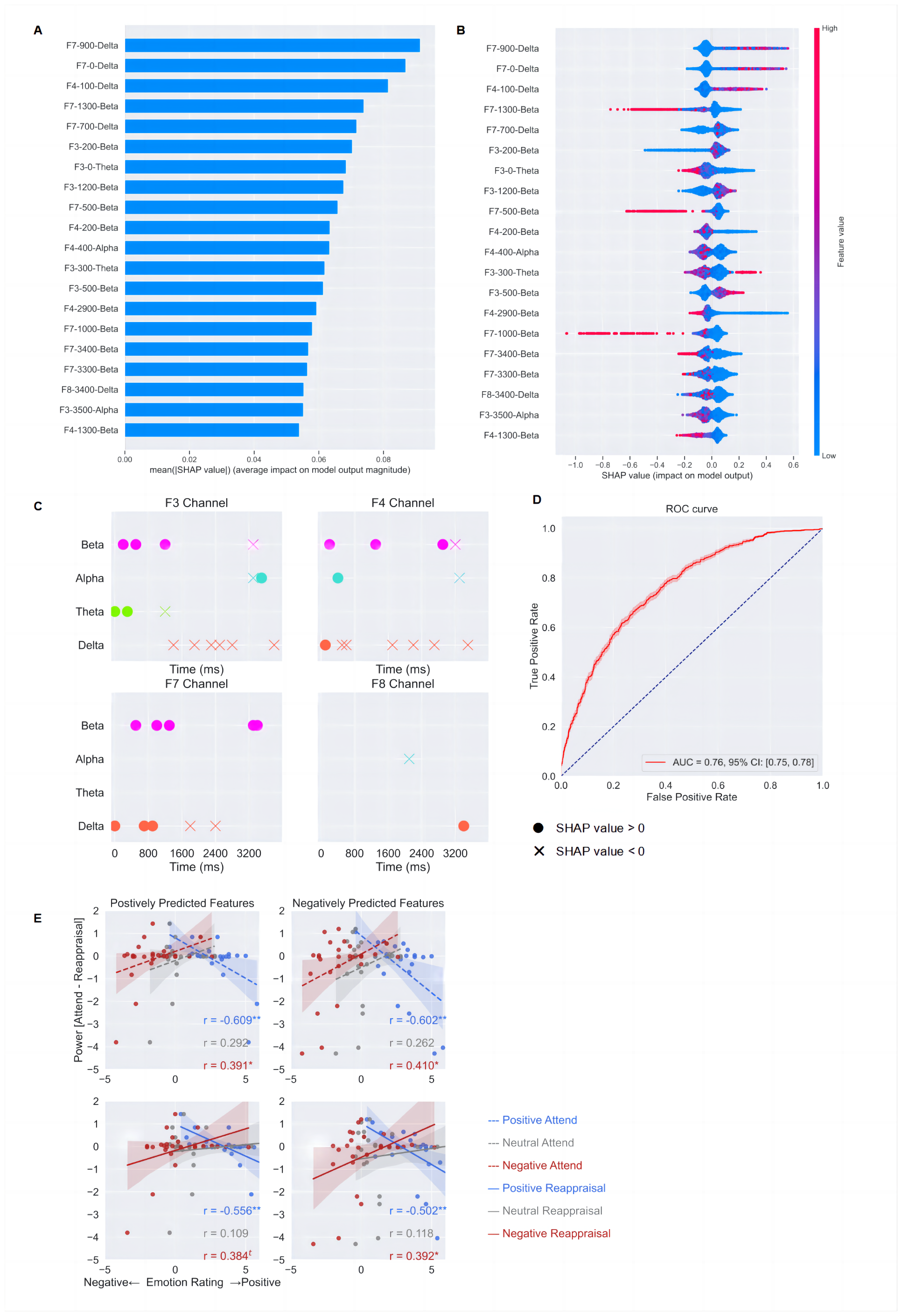
Presentation of Main Results

### Relationship Between Important Features and Emotional Ratings

In exploring the emotion regulation process, this study employed a machine learning model to predict individuals’ performance in emotion reappraisal tasks and utilized the SHAP value interpreter to analyze the model results. Furthermore, the study analyzed the correlation between the power changes (Power Attend – Power Reappraisal) of positive and negative predictive features and emotional ratings under viewing and reappraisal conditions, covering six different conditions: positive attend, neutral attend, negative attend, positive reappraisal, neutral reappraisal, and negative reappraisal. Specific results are presented in Table3. From Figure1 E, it can be observed that the energy values of features exhibit a significant correlation with emotional ratings under different emotion regulation strategies. The power changes of predictive features were significantly negatively correlated with emotional ratings during passive viewing of positive evaluations (Positive Predictive Features: *r* = -0.609, *p* = 0.001; Negative Predictive Features: *r* = -0.602, *p* = 0.001), while they were significantly positively correlated with emotional ratings during passive viewing of negative evaluations (*r* = 0.391, *p* = 0.048; *r* = 0.410, *p* = 0.038). Under cognitive reappraisal conditions, the power changes of predictive features were significantly negatively correlated with emotional ratings during positive evaluations (Positive Predictive Features: *r* = -0.556, *p* = 0.003; Negative Predictive Features: *r* = -0.502, *p* = 0.009), while they were significantly positively correlated with emotional ratings during negative evaluations (*r* = 0.384, *p* = 0.053^t^; r = 0.392, *p* = 0.047). However, no significant results were found under all neutral rating conditions.

**Table 3.** Correlation Analysis between Predictive Feature Energy Changes and Emotional Ratings under Various Conditions.

| Condition | Positive Predictive Feature |  | Negative Predictive Feature |  |
| --- | --- | --- | --- | --- |
|  | *r* | *p* | *r* | *p* |
| Positive Evaluation - Viewing | -0.609 | **0.001** | -0.602 | **0.001** |
| Neutral Evaluation - Viewing | 0.292 | 0.148 | 0.262 | 0.195 |
| Negative Evaluation - Viewing | 0.391 | **0.048** | 0.410 | **0.038** |
| Positive Evaluation - Reappraisal | -0.556 | **0.003** | -0.502 | **0.009** |
| Neutral Evaluation - Reappraisal | 0.109 | 0.596 | 0.118 | 0.568 |
| Negative Evaluation - Reappraisal | 0.384 | **0.053<sup>t</sup>** | 0.392 | **0.047** |

**BOLD:** $p < 0.05$
<sup>t</sup> marginal significant

## Discussion

Within specific EEG channels and frequency bands, brain activity at certain time points significantly influences distinguishing between different emotional or cognitive states. This study identifies critical feature points through SHAP value analysis, bearing profound implications for neuroscience and cognitive psychology. The significant brain activity changes found in specific frequency bands and time points in the LPFC area (channels F3, F4, F7, and F8) offer new insights into understanding how the brain processes emotions and performs cognitive tasks. Observing the distribution characteristics of important features (top 20 positive predictive features, top 20 negative predictive features) across channels, frequency bands, and time points leads to several insights.

First, the important positive predictive features for cognitive reappraisal predominantly emerge in the early stages of emotional experience, approximately 0-1400 ms after stimulus presentation, mainly distributed across Beta, Theta, and Delta bands. This distribution aligns with current theoretical perspectives. Cognitive reappraisal, as a cognitive control strategy, can alter an individual’s interpretation and evaluation of emotional stimuli, thereby regulating emotional responses. The early phase of emotional experience usually includes the preliminary recognition and parsing of emotional stimuli, where neural activity during this phase may be critical for successful emotion regulation. While many studies have analyzed emotional states through spontaneous EEG or traditional event-related potentials, research utilizing Event-Related Oscillations (ERO) to study emotional states is relatively less common. Emotional-related ERO research found that the Beta band is typically associated with emotion recognition (24),(25),(26),(27). Güntekin & Başar’s series of studies measured the maximum inter-peak mean amplitude of Beta (15-30Hz) from 0 to 300 ms after the appearance of emotional stimuli (emotional faces or negative pictures). Results showed that negative emotions induce greater Beta changes (bilateral electrodes in the frontal, central, and parietal lobes) compared to neutral and positive emotions. Mishra et al. (2022) discovered Beta’s functional connectivity patterns are associated with distinguishing between pleasant and unpleasant emotional stimuli(27). Theta oscillations are related to emotional expressiveness, with early Theta oscillations reflecting the initial encoding of emotionally salient sensory information (28). Research has shown that the power of Delta oscillations during psychological tasks is related to the suppression of sensory input interference. This suggests that the inhibitory Delta oscillations may regulate brain activity that should not be active during task completion (29). Frontal Delta oscillations can also reflect the level of cognitive processing in the brain’s frontal areas. According to Yener et al. (2016), the volume of the frontal lobe in elderly individuals with mild cognitive impairment positively correlates with frontal Delta oscillations (30). These results imply that frontal Delta oscillations might preliminarily process incoming emotional sensory information by pruning unnecessary cognitive activities to enhance cognitive processing efficiency. In the later part of emotion regulation, the primary negative predictive feature is the late-phase Delta band energy, possibly indicating a reduction in Delta energy during the late stages of emotion regulation. Since Delta band energy is associated with meditation and relaxation states, the reduction in Delta band energy during the later stages of emotion regulation might relate to decreased emotional responses post-regulation (see review (31)). Notably, although many studies claim Alpha oscillations play a key role in emotion regulation (see review (32)), at least in the top 20 positive and negative predictive features of this study, Alpha oscillations contribute less. One possible reason is that this study ranks all features with a 100 ms time window, making it difficult for Alpha oscillations’ SHAP values to be captured if Alpha oscillations continuously participate in the process of emotion regulation. However, some studies suggest Alpha oscillations may not significantly impact emotion regulation (33). Therefore, future research needs to improve data analysis methods to further examine the role of Alpha oscillations.

This study reveals the relationship between individual emotional ratings and specific EEG features (changes in predictive feature energy) during the emotional regulation process. The machine learning model, aided by the SHAP value interpreter, demonstrates a significant correlation between the changes in energy of predictive features and emotional ratings across different emotional tasks (viewing and reappraisal). Specifically, both positive and negative predictive features show a negative correlation with the changes in positive emotional experience (whether in passive viewing or up-regulation of emotions) and a positive correlation with changes in negative emotional experience (whether in passive viewing or down-regulation of emotions), but are uncorrelated with changes in neutral emotional experience. Specifically, during passive viewing, the greater the ΔPower [viewing - reappraisal] of individual EEG predictive features, the lower their positive emotions triggered by positive evaluations, and similarly, the lower their negative emotions triggered by negative evaluations. During cognitive reappraisal, the greater the ΔPower [viewing - reappraisal] of individual EEG predictive features, the lower their emotional ratings for down-regulated negative emotions and the lower their capacity for up-regulated positive emotions. These results suggest that the selected positive/negative predictive features may reflect an individual’s capacity for emotional perception and judgment, indicating that individuals with greater energy in these key features under viewing conditions have poorer emotional perception and judgment abilities. This aligns with the analysis that the selected important features primarily perform the identification function in the early processing of emotions. Through the application of machine learning models and SHAP value interpreters, this study reveals a significant relationship between specific EEG feature changes and emotional ratings during the emotional regulation process, offering a new perspective to understand the neural mechanisms of emotional regulation.

In terms of the application of machine learning and artificial intelligence, the discovery of these critical feature points is crucial for developing predictive models and personalized cognitive enhancement tools. Accurate temporal and frequency information can be used to train algorithms to predict individual responses to specific situations, providing support in designing personalized educational and mental health applications. Overall, these results obtained through precise temporal dynamic analysis not only deepen our understanding of the brain’s processing of complex tasks but also provide a valuable foundation for future clinical and technological applications. They reveal subtle changes in brain activity in specific cognitive and emotional tasks, offering new paths for further research, especially in precise neural mechanisms in emotional and cognitive processes. Furthermore, these findings may have significant applications in clinical practice. For instance, information on critical feature points can aid in diagnosing and treating emotional-related mental health issues, such as anxiety or depression. In mental health diagnostics, these feature points could serve as early warning signals, indicating the need for further evaluation or intervention. Similarly, in psychotherapy, this data can guide neurofeedback therapy, helping patients learn how to regulate their emotional responses, especially in training emotion reappraisal strategies.

## Conclusions

Our results highlight the significance of specific EEG features, particularly in the lateral prefrontal cortex (LPFC), in modulating emotional experiences during both passive viewing and cognitive reappraisal tasks. Notably, we observed distinct correlations between changes in EEG feature energy and emotional ratings across different emotional conditions, underscoring the role of the LPFC in early emotional processing and regulation. Furthermore, our analysis revealed the importance of temporal dynamics in emotion regulation, offering insights into the underlying neural mechanisms.

## Limitations

Despite the utility of machine learning in assisting the exploration of the temporal mechanisms involved in cognitive reappraisal processes, its interpretability remains low, and the inability to conduct causal inference warrants cautious treatment of its results. Furthermore, the discussion of mechanisms using machine learning heavily relies on feature engineering, where the selection and filtering of features could significantly alter the outcomes. Another critical limitation is the sample size, which poses a more significant constraint on EEG classification. This study included 4680 samples, but the high repeatability within subjects increases the risk of overfitting. In summary, causal evidence from laboratory studies is particularly crucial for the investigation of this topic.

Additionally, the experimental paradigm used in this study is based on a social evaluation paradigm, reflecting the processing characteristics of social negative emotions. Whether these results can be similarly interpreted within the broader domain of emotional regulation requires further validation.

## ACKNOWLEDGEMENTS

I extend our sincere gratitude to Li Sijin for providing the data used in this study.

